# Mixed fatty acid-phospholipid protocell networks

**DOI:** 10.1101/2021.03.08.434432

**Authors:** Inga Põldsalu, Elif Senem Köksal, Irep Gözen

## Abstract

Self-assembled membranes composed of both fatty acids and phospholipids are both permeable for solutes and structurally stable, which was likely an advantageous combination for the development of primitive cells on the early Earth. Here we report on the solid surface-assisted formation of primitive mixed-surfactant membrane compartments, *i.e.* model protocells, from multilamellar lipid reservoirs composed of different ratios of fatty acids and phospholipids. Similar to the previously discovered enhancement of model protocell formation on solid substrates, we achieve spontaneous multi-step self-transformation of mixed surfactant reservoirs into closed surfactant containers, interconnected via nanotube networks. Some of the fatty acid containing compartments in the networks exhibit colony-like growth. We demonstrate that the compartments generated from fatty acid-containing phospholipid membranes feature increased permeability coefficients for molecules in the ambient solution, for fluorescein up to 7*10^-6^ cm/s and for RNA up to 3.5*10^-6^ cm/s. Our findings indicate that surface-assisted autonomous protocell formation and development, starting from mixed amphiphiles, is a plausible scenario for the early stages of the emergence of primitive cells.

## Introduction

Contemporary living cells are thought to have originated from primitive protocells. A widely accepted protocell model is a spherical compartment, self-assembled from structurally simple amphiphiles freely suspending in an aqueous “soup”. The formation of closed compartments enveloped by a semi-permeable surfactant membrane would have allowed the segregation and confinement of prebiotic chemical compounds, which provided protection from dilution and prevented interference from other compouxnds in the external environment.

We had recently reported a solid-surface mediated pathway of autonomous protocell formation which results in formation of phospholipid compartments, physically connected to each other via a network of lipid nanotubes^1^. The compartments were shown to draw the energy necessary for the shape transformation of their membranous precursor structures from the surface. During maturation they encapsulate ambient constituents and separate from the network with a gentle hydrodynamic flow, which provides a feasible means to migrate to remote locations. The reported mechanism can, with a minimal set of assumptions, explain protocell self-formation and development, offer a primitive pathway to replication, and has enabled us to formulate a new division hypothesis^2^. The protocell models reported in this study consisted purely of phospholipids (PLs). Several earlier studies have provided evidence that phospholipids (PLs) involved in the assembly of the first protocells could have been synthesized under prebiotic conditions^3–5^.

Another lipid species, the fatty acids (FAs), are structurally simpler than phospholipids, can provide key advantages to protocell membranes, including high permeability to ions and small charged molecules, and enable continuous exchange of lipids which promotes vesicle growth^6, 7^. These characteristics make FA-rich membranes a promising candidate for the first protocells, adding favorably to the structural stability and longevity that purely phospholipid-based membranes provide. FAs might have been synthesized earlier under prebiotic conditions^8, 9^ or have possibly arrived in our geosphere via meteorite impacts^10, 11^ where under favorable conditions they might have formed micelles and early prebiotic compartments. However, the presence of high concentrations of divalent cations or peptide sequences, which are necessary for nonenzymatic replication of RNA, can disrupt fatty acid membranes^12^. One way for the single chain lipids, *e.g.* fatty acids, to become more stable is via high favorable interactions with other membrane amphiphiles. Di-acyl lipid mixtures have been shown to enable vesicle growth and stability in aqueous environments^13^, also in the presence of divalent cations^6^.

Here we report the surface-assisted formation of compartments consisting of PLs with an increasing fraction of FAs. Upon contact with a SiO2 surface, bulk reservoirs of mixed amphiphiles act in a manner similar to the ones made purely from phospholipids, and spontaneously transform into protocell- nanotube networks. The model protocells with fatty acid-containing membranes appear to promote the colony-like growth of multiple adjacent compartments. We demonstrate that the fatty acid-containing protocells take up ambient constituents such as fluorescein and RNA. We show compelling evidence that primitive cells could have self-assembled from mixed amphiphiles on solid supports, and exhibit collective growth. Compared to pure phospholipid model protocells, the mixed membranes feature higher permeability for ambient constituents while remaining stable for hours up to days.

## Results

When multilamellar vesicles (MLVs), composed solely of phospholipids, come in contact with solid SiO_2_ substrates, a series of topological transformations occur^1^ (**Fig. 1a-e**). The sequence depicted there has been demonstrated to be a robust pathway for the autonomous transformation of pure phospholipid membranes on flat silicon dioxide surfaces.

**Figure 1.**
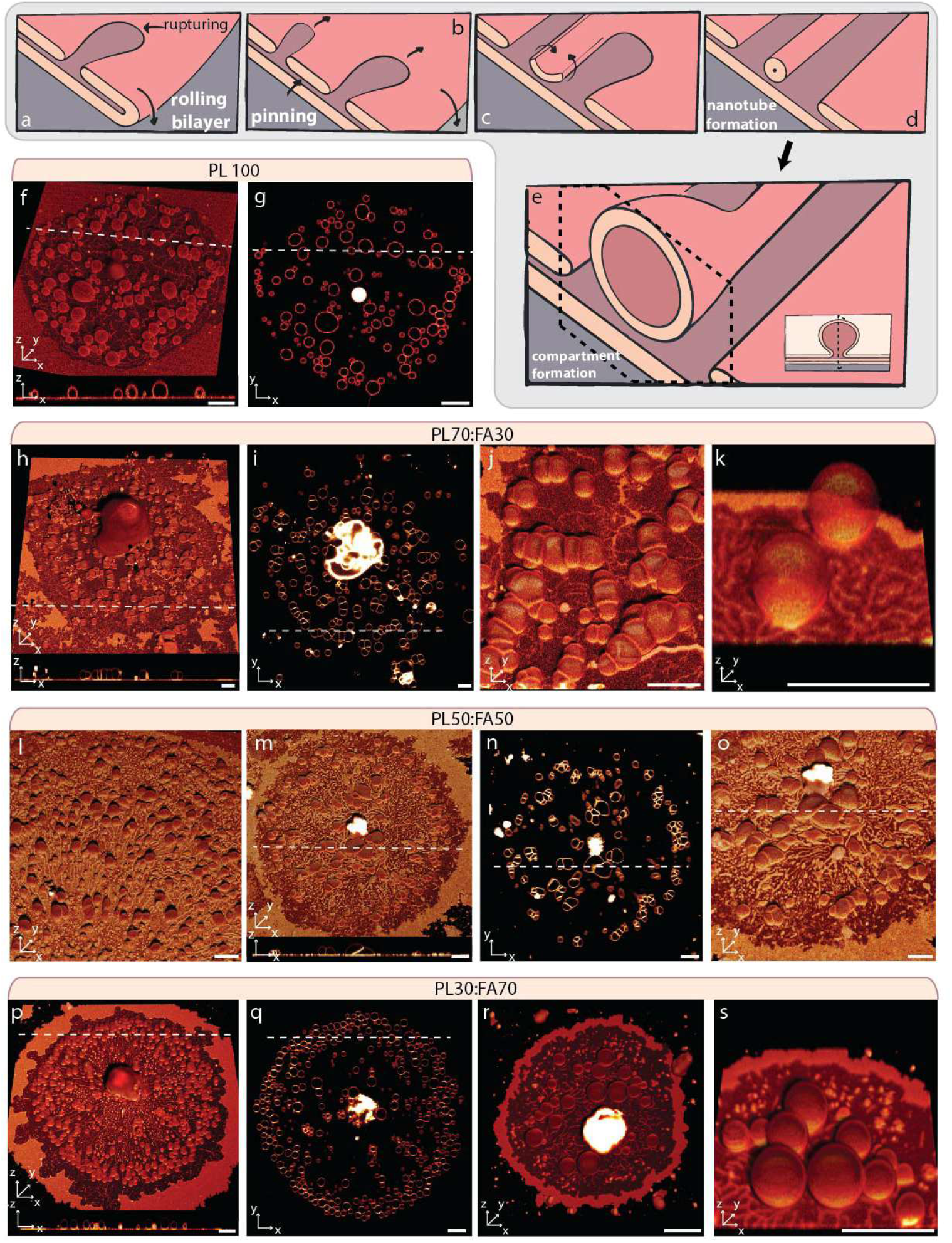
Formation of model protocells from lipid membranes containing phospholipid and fatty acid mixtures. **(a-e)** Schematic drawing, showing the topological membrane transformations after contact of a surfactant reservoir with the solid surface. **(a)** The membrane spreads as a double bilayer membrane, exhibiting rolling bilayer motion. The distal membrane eventually ruptures due to tensile stress. **(b)** While ruptures propagate, some membrane regions maintain due to inter-bilayer pinning. **(c)** Elongating membrane threads are produced, which form nanotubes **(d)** to alleviate the resulting high edge tension. **(e)** Fragments of the nanotubes swell and form spherical lipid compartments. **(f-s)** Confocal micrographs showing model protocells originating from lipid reservoirs consisting of: (f-g) 100% PL, (h-k) 70PL:30FA, (l-o) 50PL:50FA, (p-s) 30PL:70FA. (f, h, j, k, l, m, o, p, r, s) 3D micrographs, (g-i-n-q) 2D micrographs of (f-h-m-p). The cross-sectional side views along the white dashed line in (f-h-m-p) is presented at the bottom each panel. Scale bar: 10 μm.

The initial preparation step is the spreading of the multilamellar lipid reservoir in form of a circular double lipid bilayer membrane^14^. During this wetting process the proximal membrane (lower membrane with respect to the surface) continuously adheres onto the solid substrate and drags the distal (upper) membrane along, resulting in rolling bilayer motion^15^. The rolling continues until the membrane tension reaches lysis tension (5-10 mN/m^16^) and distal bilayer ruptures. **Fig. 1a** shows a schematic drawing depicting the cross section of the spreading edge of a rupturing double bilayer membrane. While the lipid material is displaced due to rupturing, some fractions of the distal membrane are immobilized due to the pinning between the two lipid bilayers (**Fig. 1b**). The pinning is caused by the fusogenic ions in the ambient buffer *e.g.* Ca^2+^^14, 17^. The rupture propagation causes the elongation of the membrane regions between the pinned sites and the displacing membrane edge due to rupturing (**Fig. 1c**). Elongated membrane edges lead to the increasing cost of line energy *E = γl,* where *γ* is edge tension (5-10 pN) and *l* is the length of the elongated membrane edge. To reduce this cost, the lengthy membrane regions wrap up forming lipid nanotubes (**Fig. 1d**). Fragments of these nanotubes over time swell into giant unilamellar compartments (**Fig. 1e**). **Fig. 1f** shows a 3D confocal micrograph of a lipid patch originating from MLVs consisting 100% of phospholipids (control). The cross-sectional profile view (x-z plane) along the white dashed line has been presented below the 3D micrograph (valid also for panels **h, m** and **p**). **Fig. 1g** shows a 2D cross sectional view of the region in panel (**f**) in x-y plane.

**Fig. 1 h-s** shows lipid compartments originating from multilamellar reservoirs containing a mixture of PLs and FAs and formed as a result of the process described above: (**h-k**), 30% FA, (**l-o**), 50% FA and (**p-s**) 70% FA. A full list of the prepared compositions as well as additional confocal micrographs is provided as supporting information (**SI 1-2**). We note that compartments also form from reservoirs consisting of nearly pure FAs (99% FA with 1% fluorophore-conjugated PL for visualization) under identical conditions (**SI 3**), which we did not include in the current investigation.

The model protocells formed from mixed FA-PL containing membranes generally resemble in properties and behavior those obtained from pure phospholipid mixtures. One prominent difference is the shape morphologies of FA containing compartments, which grow collectively in a colony-like manner with their adjacent bilayers in contact, and adopt peculiar morphologies (**Fig. 1i-j** and **m-n,** *cf.* SI 2 for examples regarding 70FA:30PL). Protocell colonies have been previously hypothesized^18^ and artificially assembled with acoustic wave patterning^19^, by adding Mg^2+^ as high as 50 mM to initially isolated compartments^6^, or by supplying polymers such as poly-L-lysine (PLL)^20^. To the best of our knowledge, our results constitute the first example of model protocell colonies that emerge and grow in clusters without any directed agitation or addition of supporting molecules or ions to the experimental environment.

To provide a means of comparison, we analyzed the shape of a total of 2155 model protocells of all compositions, and determined a roundness factor (**SI 4**). Most of the 100% PL containing compartments have nearly perfectly circular cross sections, while the shapes of the fatty acid containing membranes vary, some with roundness values as low as 0.2. If the FA containing compartments are isolated, they adopt circular shapes (**Fig. 1k**). The mutual adhesion of giant unilamellar vesicles (GUVs) was previously observed in the presence of mM concentrations of Ca^2+^^21^ and adhesion energies were calculated from the contact angles between the two GUVs^22^. The bilayer-to-bilayer adhesion energies in 1-10 mM Ca^2+^ were determined to be 0.02-0.03 mN/m. This is well below the lysis tension of the membrane (5 mN/m^16^), therefore the adjacent colony-like protocells are able to remain stable for long periods of time (days).

In order to understand if the FAs were properly assembled into the protocell membranes, we quantified free FA species in solution for each FA-containing sample (**SI 5**). In general, with increasing amounts of FAs contained in the prepared lipid suspensions, we determined that a decreasing amount of free fatty acids in the external solution. This is evidence that the FAs have a tendency to be incorporated efficiently into the lipid membrane. We consider this a rather surprising finding, as a previous report^6^ shows that the mixtures of increasing amount of fatty acid in phospholipid membranes leads to increasing free fatty acid concentrations in the solution. This disagreement remains unexplained, but may be due to a special interaction between the lipid species used in the study, oleic acid and phosphatidyl choline.

We investigated the membrane permeability of the surface adhered model protocells to the compounds in the ambient solution (**Fig. 2**). Briefly, with a microfluidic pipette, the vesicular compartments (**Fig. 2a**) were locally superfused with fluorescein- or fluorescently labeled RNA-containing solutions (**Fig. 2b**), resulting in the spontaneous encapsulation of these components (**Fig. 2c**). A sample time series of this experiment is shown in **Fig. 2d-g**. **Fig. 2e-g**, which is the magnified version of the region shown in a white dashed frame in **Fig. 2d**, shows confocal micrographs of the process corresponding to **Fig. 2a-c**. We performed this experiment on lipid compartments consisting purely of PLs, as well as on those consisting of PLs and FAs. We monitored the uptake, represented by the increase of the fluorescence signal inside the compartments, with laser scanning confocal microscopy.

**Figure 2.**
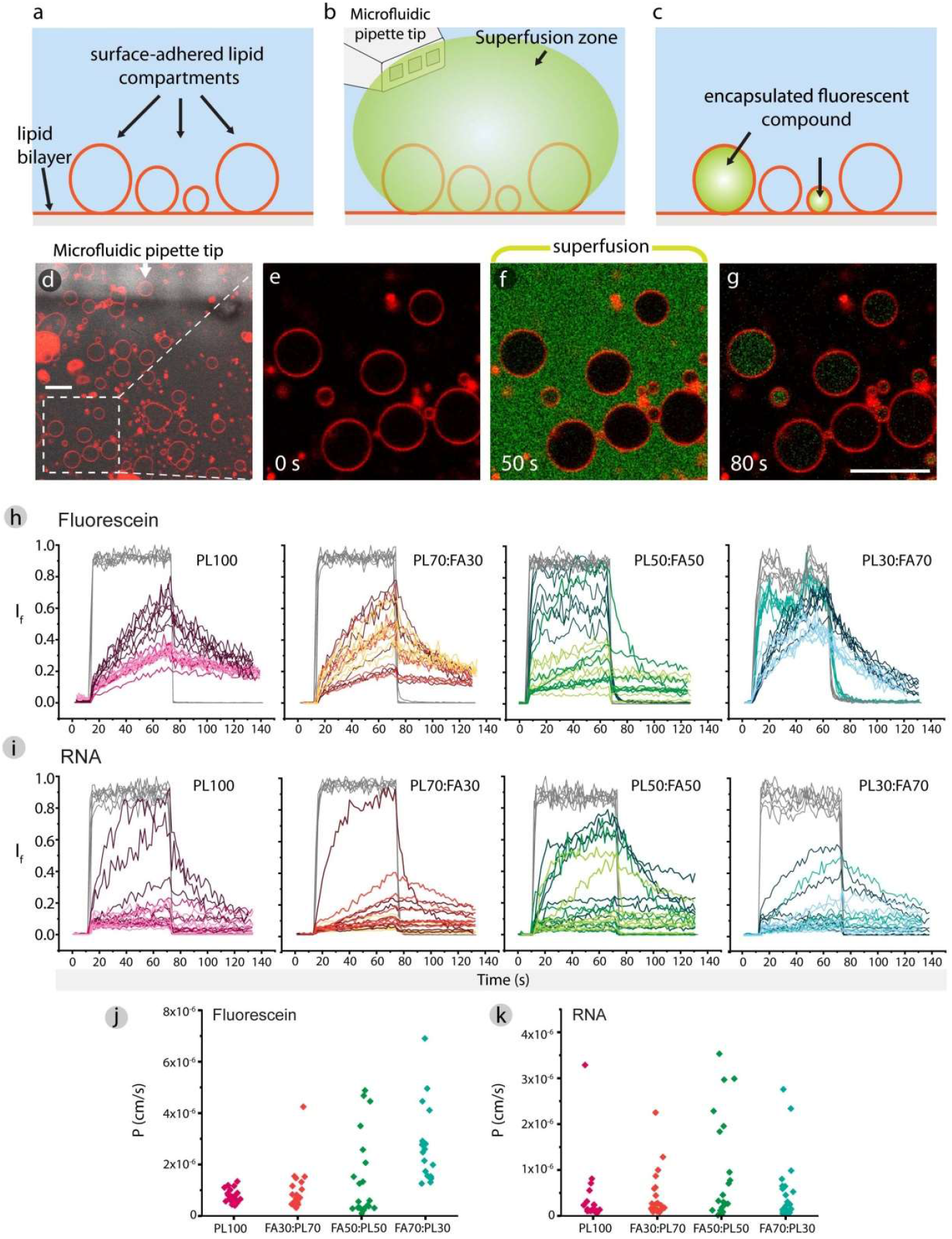
Uptake of fluorescein and RNA by model protocells from phospholipid-fatty acid membranes. **(a-c)** Schematic drawing summarizing the experiment. Adhered model protocells (a) were superfused with the compound of interest for 1 min resulting in the spontaneous encapsulation of the compound by some of the compartments. **(e-g)** Confocal micrographs corresponding to (a-c). (e-g) are magnified version of the region in **(d)** framed with white dashed lines. **(h)** Uptake of fluorescein, and **(i)** RNA, by model protocells. Permeability coefficients for **(j)** fluorescein and **(k)** RNA calculated for each vesicle in (h) and (i), respectively. Scale bar: 10 μm.

The uptake of fluorescein by lipid compartments with varying fatty acid composition is shown in **Fig. 2h** and the uptake of the fluorescently labeled RNA in **Fig. 2i**. Each plot shows the fluorescence intensity in a region of interest (ROI) during and after superfusion. Every colored plot represents a ROI positioned inside a compartment and different shades in the same graph represent different experiments. The ROIs positioned outside the compartments are represented in gray color. Confocal micrographs showing all ROIs are provided in the supporting material (**SI 6**).

Using the data presented in **Fig. 2h-i** we calculated the permeability coefficient *P* for every compartment^23^. **Fig. 2j-k** shows *P* for fluorescein and RNA, respectively, grouped for each membrane composition. For fluorescein, *P* is consistent among the 100% PL protocells and is ~0.5-1*10^-6^ cm/s. These values are in good agreement with the ones reported by Mohanan, *et al*.^24^ on the uptake of similarly sized drug molecules by bacteria-derived phospholipid vesicles. The *P* values increase with increasing fatty acid concentration and are spread out from very low to as high as 7*10^-6^ cm/s. We have previously shown the encapsulation of fluorescein by surface adhered phospholipid compartments^1, 25^ where we proposed the uptake to be via nano-sized transient pores and predicted the quantity of uptake with a finite element method (FEM)-based model^25^. The uptake rates presented in **Fig. 2h** for the PL protocells agrees well with this prior work. For the FA-containing protocells, some of the uptake can result from the formation of such transient openings due to mechanical disturbance during superfusion, but this alone cannot explain the increasing permeability coefficients with increasing fatty acid concentrations. We therefore assume that the higher *P* values obtained upon increased FA content is due to the increased disorder caused by the incorporation of FAs to the membrane interior^26^. Membrane perturbing effects of FAs and their ability to reduce the membrane permeability barrier are well-established^26, 27^.

For RNA, the uptake behavior based on the fatty acid content is similar, but the permeability values are generally lower, the majority of the sample being below 2*10^-6^ cm/s, compared to those for fluorescein. The lower permeability to RNA is plausible, as the fluorophore-conjugated RNA is significantly larger in size than the fluorescein molecule (M_w_ of fluorescein: 332.31 g/mol; M_w_ of fluorophore conjugated RNA: 3767.60 g/mol). The compartments with the highest permeability values for RNA are those containing 50% FA. It has been previously hypothesized that the RNA permeation exhibited by protocellular membranes occurs through formation of supramolecular RNA complexes, which adhere to and destabilize the lipid membranes, leading to transient openings by action from one side^28^. The time scale of this disruption is relatively slow; it measures in hours^29, 30^ compared to the uptake on the order of seconds in our experiments. We therefore think that the RNA uptake by the model protocells in our system is occurring without the formation of any RNA complexes, but is directly due to the intrinsic membrane perturbing mechanisms attributed to the FAs as described above.

## Conclusion

We are able to show that solid surface-assisted autonomous formation of model protocells is not limited to pure phospholipids, but extends to membranes containing different amounts of fatty acids. Furthermore, the resulting protocells are in a similar fashion physically connected via networks of nanotubes, some exhibiting colony-like growth. In our study, represented by the model solutes fluorescein and 10 base RNA, we demonstrated that the protocells take up and maintain constituents of the surrounding aqueous environment. The protocell membranes have generally increasing permeability with increasing fatty acid concentrations. Our findings indicate that the surface-assisted protocell formation and development of mixed amphiphiles is a plausible scenario for the early stages of the emergence of primitive cells. We conclude that protocell nanotube network structures composed of different lipid species would be a novel framework of structures to accommodate and study prebiotic early Earth chemistry.

## Materials and Methods

### Preparation of lipid suspensions

Various compositions of lipid mixtures were prepared using the dehydration-rehydration method described by Karlsson *et al*.^31^. A full list of the mixtures has been provided in **SI 1**. Briefly: fatty acids, phospholipids (all lipid products from Avanti Polar Lipids, USA) and phospholipid conjugated fluorophore 16:0 Liss Rhodamine PE (Avanti Polar Lipids, USA) were dissolved in chloroform (Merck Life Science, Norway) and methanol (VWR International) 2:1 mixture to a final concentration of 10 mg/mL. The solvent was evaporated in a rotary evaporator (Büchi, Switzerland) under reduced pressure (20 kPa) over 6 h. Subsequently, the dry lipid film was rehydrated with 3 mL of 0.2 M 8.5 pH Bicine (Merck Life Science, Norway) buffer (pH adjusted with NaOH (Merck Life Science, Norway)) and 30 μL of Glycerol (Merck Life Science, Norway). The rehydrated lipids were kept overnight at 4 °C and thereafter swirled until there was no dry lipid residue. If swirling was not sufficient, the lipids were sonicated for 5 s.

### Preparation of observation chamber

4 μL of lipid suspension was placed on a coverslip (MEZ102460, Menzel Gläser, Germany) and dehydrated in an evacuated desiccator for 20 min. The dry lipid film was dehydrated with 500 μL of 10 mM HEPES (Merck Life Science, Norway) and 100 mM NaCl (pH 7.8 adjusted with NaOH) buffer for 10 min. The suspension was transferred into an open-top observation chamber (15 x 15 x 0.5 mm) with a SiO2 substrate containing 1 mL of 10 mM HEPES, 100 mM NaCl (Merck Life Science, Norway) and 4 mM of CaCl_2_ (Merck Life Science, Norway) (pH 7.8 adjusted with NaOH).

### Surface Fabrication

Substrates were fabricated by depositing 84 nm of SiO2 on a glass coverslip (MEZ102460, Menzel Gläser, Germany) using electron-beam physical vapor deposition with the Angstrom - E-beam and thermal PVD EvoVac (Ångstrom Engineering, Finland). Before deposition of the organic material, the surfaces were plasma treated (Diener electronic GmbH & Co KG, Germany) for 5 min.

### Microscopy imaging

A confocal laser scanning microscopy system (Leica SP8, Germany), with HCX PL APO CS 40x oil objective NA 1.3 was used for acquisition of confocal fluorescence micrographs. The excitation/emission wavelengths for Rhodamine was 560/583, and for fluorescein, 488/508.

### Encapsulation of dyes and RNA

To expose the matured surface-adhered protocells to the encapsulation media, an open-volume microfluidic pipette (Fluicell AB, Sweden) was used together with 3-axis water hydraulic micromanipulator (Narishige, Japan). The encapsulation media consisted of 10 mM HEPES (Merck Life Science, Norway), 100 mM NaCl (Merck Life Science, Norway), 4 mM CaCl_2_ (Merck Life Science, Norway) and 25 μM fluorescein sodium salt (Merck Life Science, Norway), or 25 μM Fl-conjugated 10 base-long Poly-Adenine RNA oligonucleotides (Dharmacon, USA), pH 7.8.

### Image processing/Analysis

3D fluorescence images were reconstructed using the Leica Application Suite X Software (Leica Microsystems, Germany). The colors assigned to the labelled lipid membranes are resulting from the false coloring of the grayscale images of the fluorescence signals and are assigned arbitrarily. Image enhancements for display figures and videos were performed with the NIH Image-J and Adobe Photoshop CS4 (Adobe systems, USA). Schematic drawings and image overlays were created with Adobe Illustrator CS4 (Adobe Systems, USA). Fluorescence intensity profiles were obtained with using NIH Image-J and after applying median filtering to the regions of interest, plotted in Origin 2020 (OriginLab Corporation, USA). For roundness analysis, background removal was done with Adobe Photoshop CS4 (Adobe Systems, USA) and particle analyses were performed with the NIH Image-J Software. 3D histograms were plotted with Matlab R2018a. Roundness is defined as 4*area/(π*major_axis^2). Permeability (P) was calculated according to Cama *et al.^23^* using the equation:

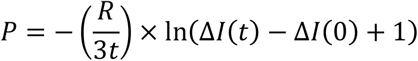

*R* is the vesicle radius, *t* the time taken for the vesicle to move from the initial (t=0) to the final (t) detection point, and *ΔI* is the normalized fluorescence intensity difference between the interior and the exterior of the 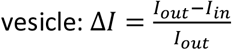.

## Acknowledgements

This work was made possible through financial support obtained from the Research Council of Norway (Forskningsrådet) Project Grant 274433, UiO: Life Sciences Convergence Environment, start-up funding provided by the Centre for Molecular Medicine Norway & Faculty of Mathematics and Natural Sciences at the University of Oslo. I.P. greatly acknowledges the European Union’s Horizon 2020 research and innovation programme under the Marie Skłodowska-Curie grant agreement No 801133.

## Author contributions

I.P., E.K. performed research and analyzed data, I.G. suggested the investigation of surface-assisted formation with fatty acid containing membranes, designed the research and supervised the project. I.P., E.K. and I.G. wrote the paper.

