## Supporting Information for "Mixed fatty acid-phospholipid protocell networks"

**This PDF file includes:**

Supplementary text

S1. Lipid compositions

Table S1

S2. Protocell formation from phospholipid-fatty acid mixtures

S3. Protocell formation from 99% fatty acids

S4. Roundness analyses

S5. Free fatty acid quantification

S6. Encapsulation of fluorescein and RNA inside the protocells (ROIs)

S7. Supplementary movies

Captions for movies S1

References for SI

**Other supplementary materials for this manuscript include the following:**

Movies S1

**S1. Lipid compositions**

**Table 1.** A full list of the lipid mixtures.

| Surface | Lipid Composition | *PL:FA ratio (wt%)* | Fluorophore (1%) | Vesicle Formation |
|---|---|---|---|---|
| $SiO_2$ | S:E | **50:49** | 16:0 Liss Rhod PE | + |
| | S:M | **70:29** | 16:0 Liss Rhod PE | + |
| | S:M | **50:49** | 16:0 Liss Rhod PE | + |
| | S:M | **29:70** | 16:0 Liss Rhod PE | + |
| | S:M | **9:90** | 16:0 Liss Rhod PE | + |
| | S:O | **49:50** | 16:0 Liss Rhod PE | + |
| | S:O | **29:70** | 16:0 Liss Rhod PE | + |
| | S:O | **9:90** | 16:0 Liss Rhod PE | rare |
| | PE:PG:CA:M | **33:14:3:49** | 16:0 Liss Rhod PE | + |
| | E:M | **69:30** | 16:0 Liss Rhod PE | + |
| | E:M | **49:50** | 16:0 Liss Rhod PE | + |
| | E:M | **29:70** | 16:0 Liss Rhod PE | + |
| | E:M | **9:90** | 16:0 Liss Rhod PE | rare |
| | M | **99** | 16:0 Liss Rhod PE | rare |
| | O | **99** | 16:0 Liss Rhod PE | rare |
| | P | **99** | 16:0 Liss Rhod PE | rare |
| | DA | **99** | 16:0 Liss Rhod PE | rare |

***Abbreviations used in the table:***
***S*** *: Soybean polar lipid extract*
***E*** *: E.coli polar lipid extract*
***PE*** *: L-α-phosphoethanolamine (E.coli)*
***PG*** *: L-α –phosphatidylglycerol (E.coli)*
***CA*** *: Cardiolipin (E.coli)*
***M*** *: Myristoleic acid*
***O*** *: Oleic acid*
***P*** *: Palmitoleic acid*
***DA*** *: Decanoic acid*
***16:0 Liss Rhod PE*** *: 16:0 Liss Rhodamine PE*

**S2. Protocell formation with phospholipid-fatty acid mixtures**

The 3D confocal micrographs presented below show protocell-nanotube networks autonomously formed on solid substrates from various PL-FA mixtures. Compositions of the lipid mixtures for each figure set are shown in the upper left corner of the images. The abbreviation used are listed in **S1**.

**E69:M30 >**

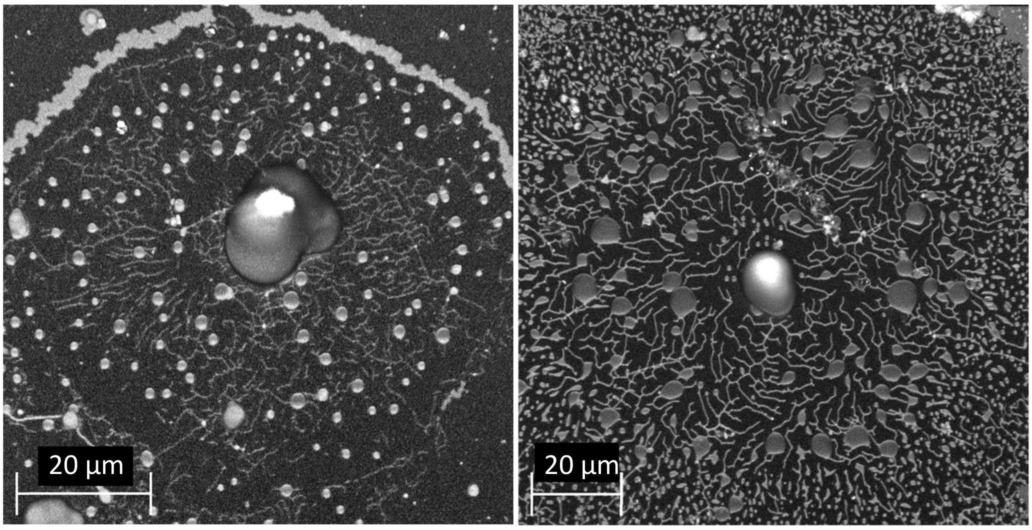


**S69:M30 >**

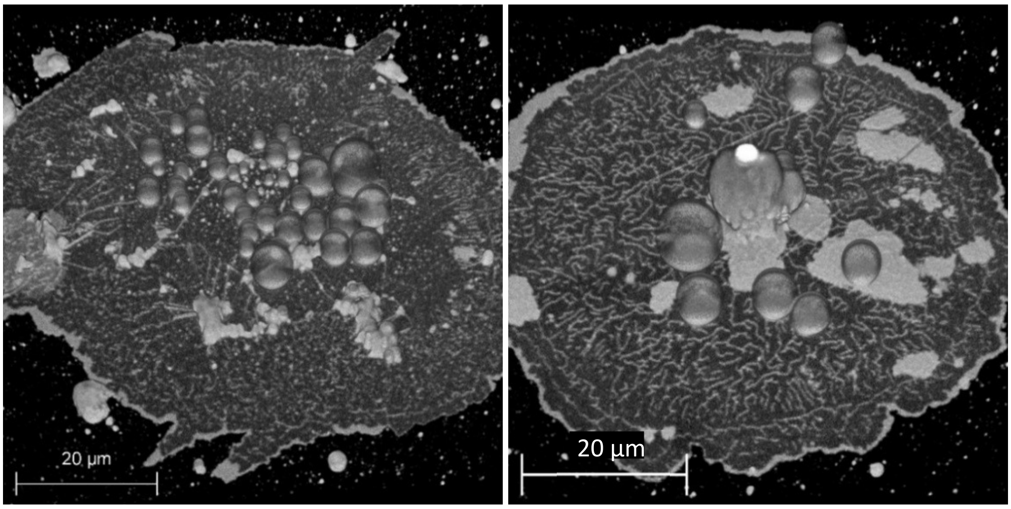


**E49:M50 >**

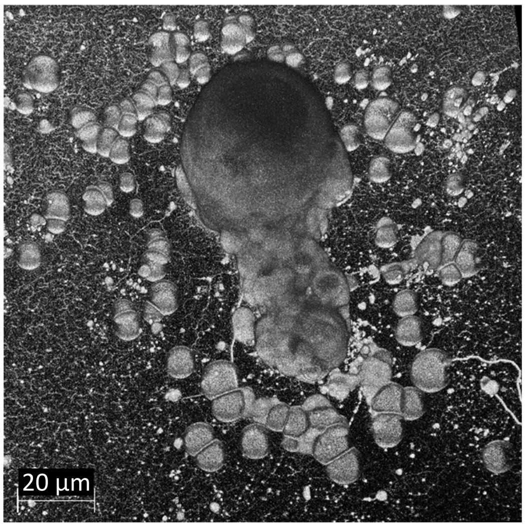

**S49:M50 >**

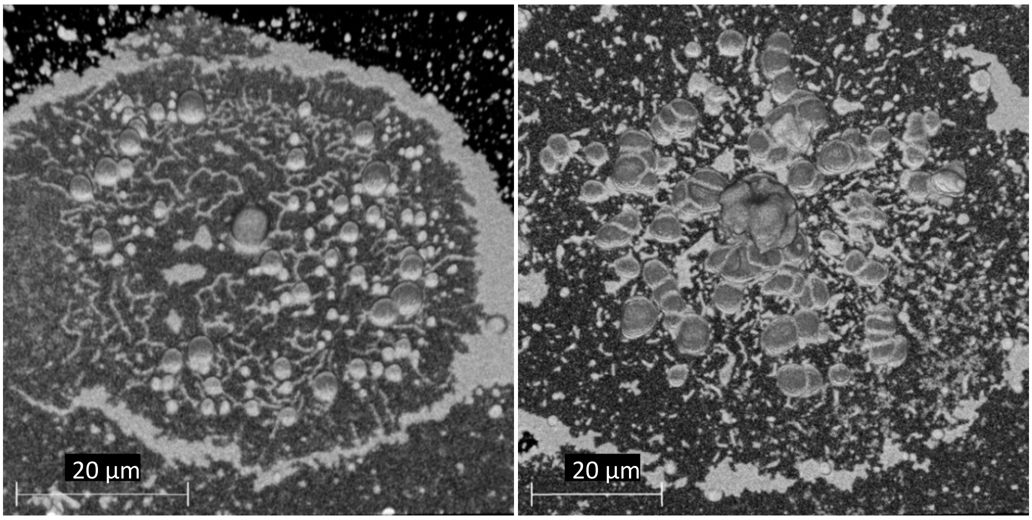


**S49:O50 >**

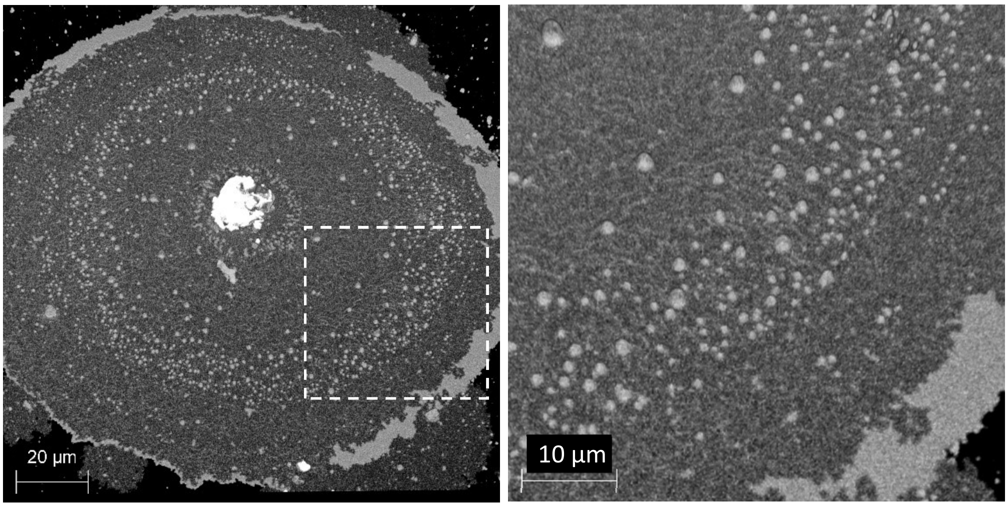


**E29:M70 >**

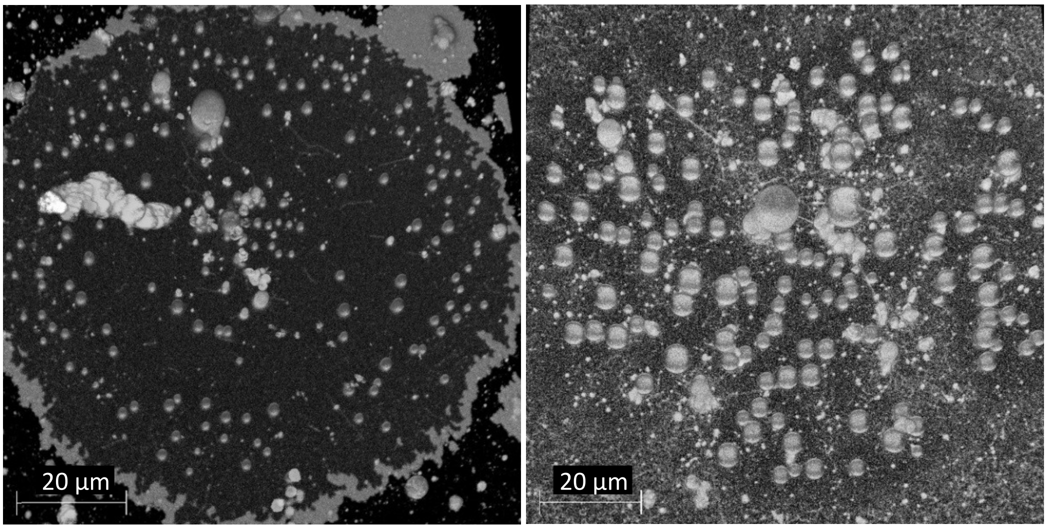

**S29:M70 >**

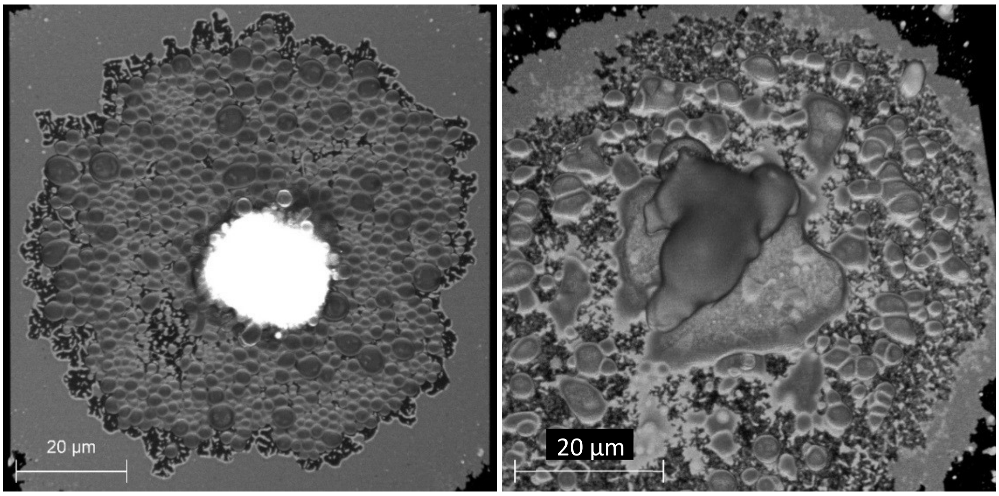


**S29:O70 >**

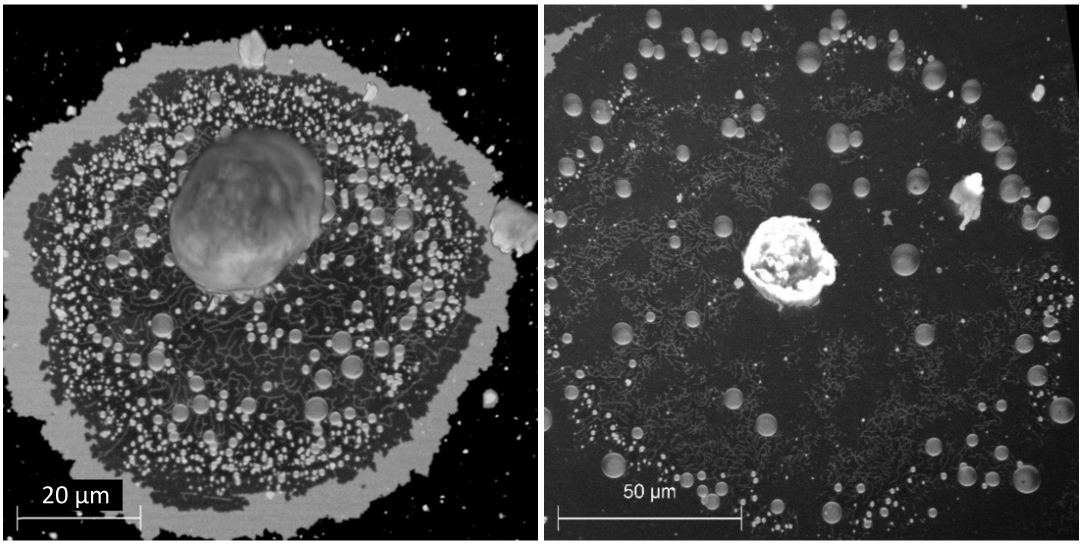


**E9:M90 >**

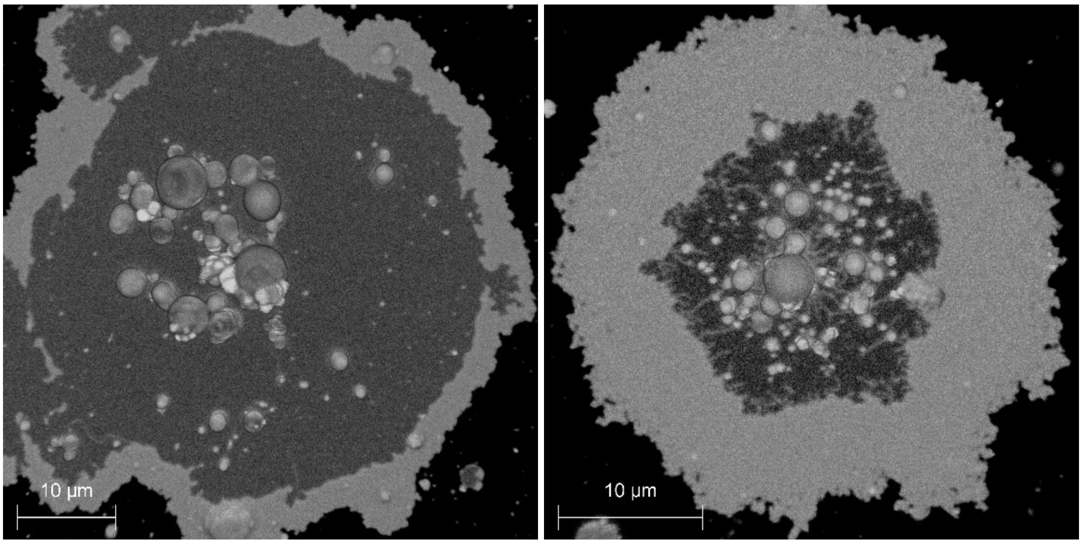

**S9:M90 >**

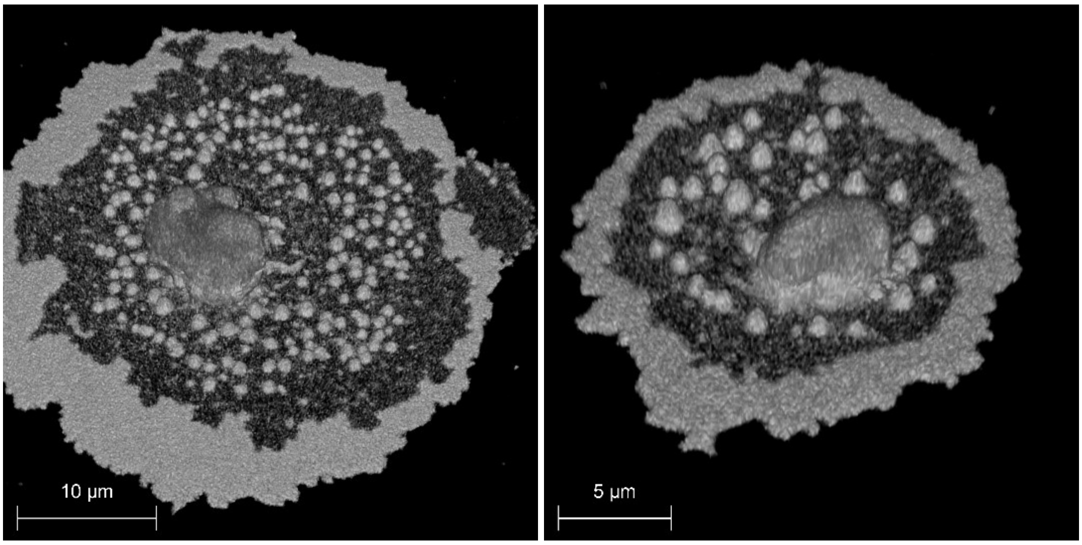


**S9:O90 >**

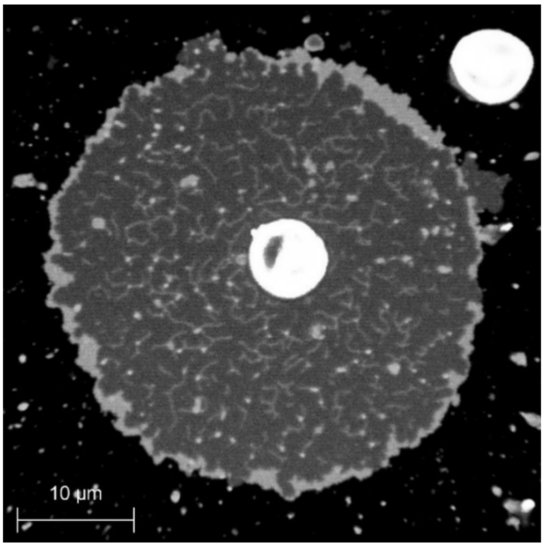

**S3. Protocell formation from 99% fatty acids**

3D-reconstructed confocal micrographs below show the formation of model protocells from 99% fatty acids. We observed nanotube-mediated protocell formation from several types of fatty acids. The comparatively low yield of pure FA structures, compared to the PL-containing mixtures, calls for a separate investigation. The current findings suggest that optimization of external parameters, such as ion strength, solution composition, temperature, and pH, might reveal conditions under which pure FA compartment networks are formed in high yield, such that permeability and stability studies can be performed on a sufficiently large vesicle population. C*f.* **S1** for abbreviations.

**P99 >**

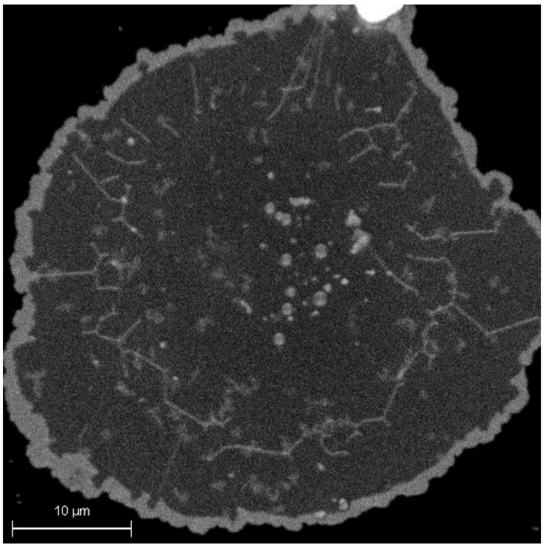

**DA99 >**

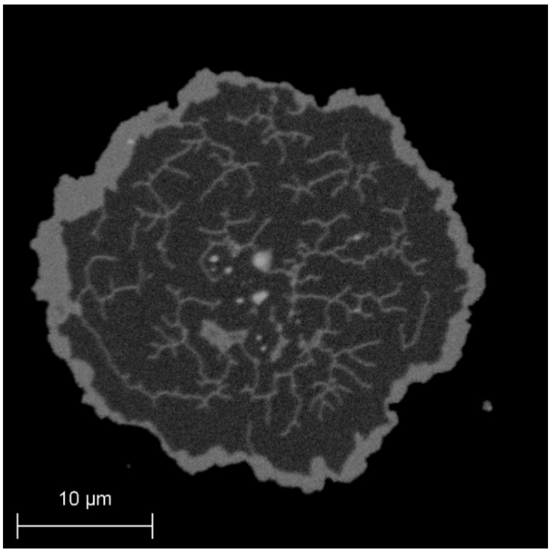


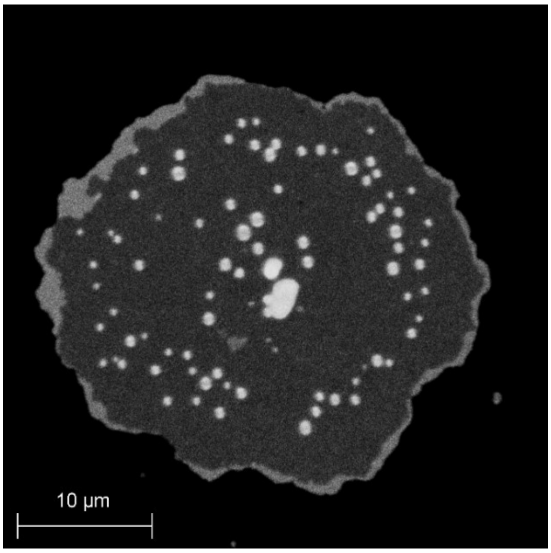


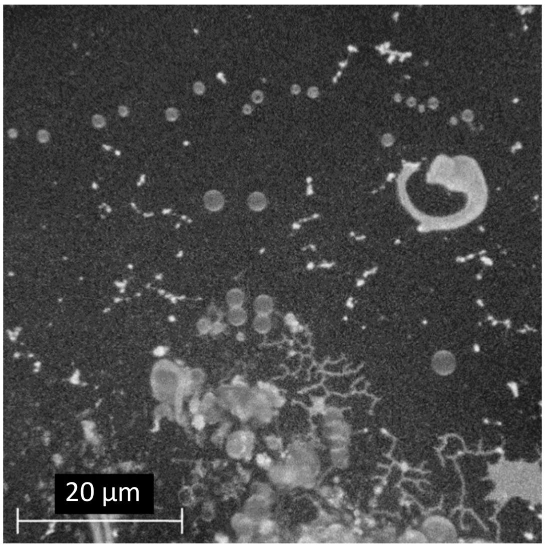


**O99 >**

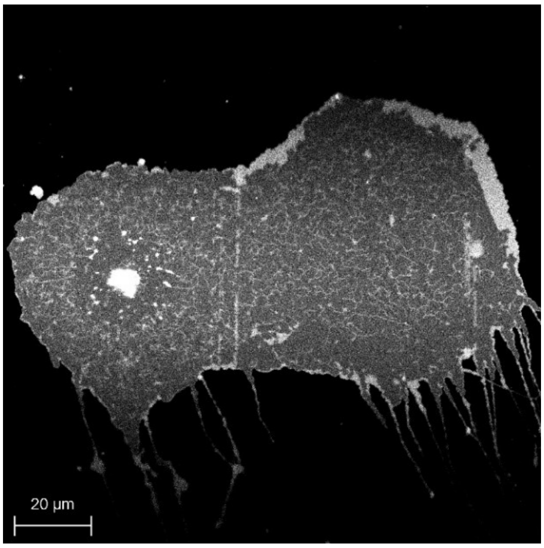

**M99 >**

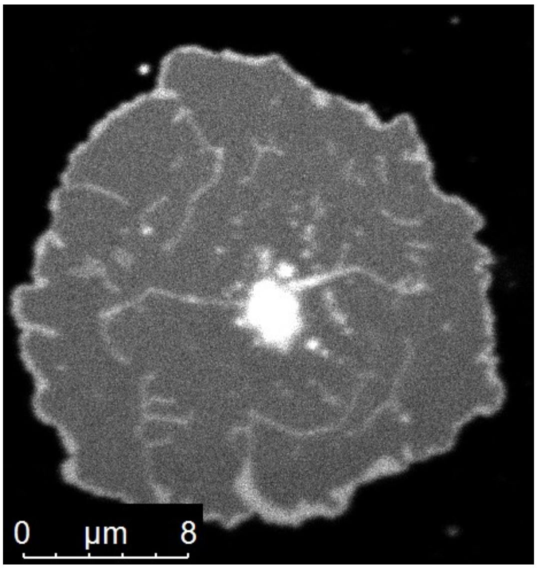


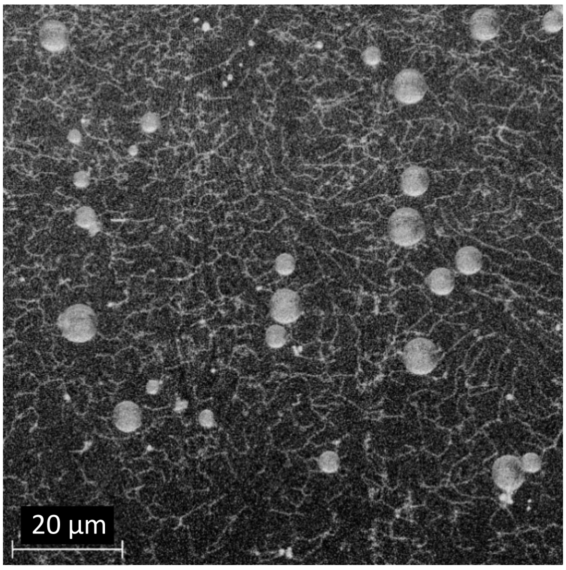


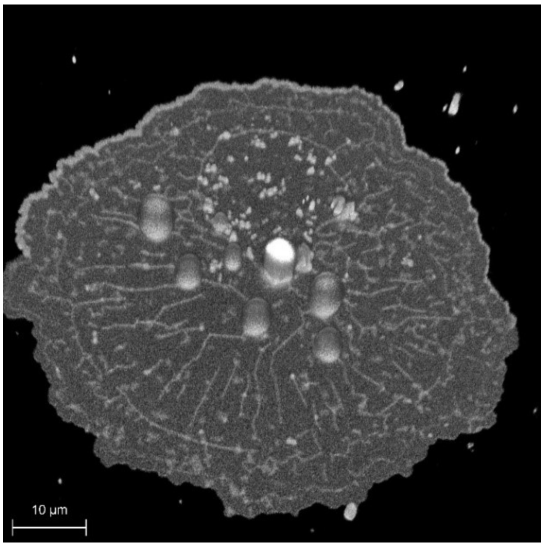


**S4. Roundness analyses**

Roundness analyses were performed to characterize the different morphologies observed in FA containing protocells. After removal of the image background with Adobe Photoshop CS4 (Adobe Systems, USA), particle analyses were performed with the NIH Image-J Software. Roundness is defined as 4*area/(π*major_axis^2). The results of the calculations for the roundness of lipid compartments formed from two different PL containing mixtures; S:M (**Figure S4a**) and E:M (**Figure S4b**) are presented below. The histograms were plotted in Matlab R2018a. The figures above each histogram indicate the number of compartments analyzed for that particular mixture. The analyses of 290 model protocells made from 100% phospholipids containing S50:E50 are shown both in (a) and (b). *Cf.* **S1** for abbreviations.

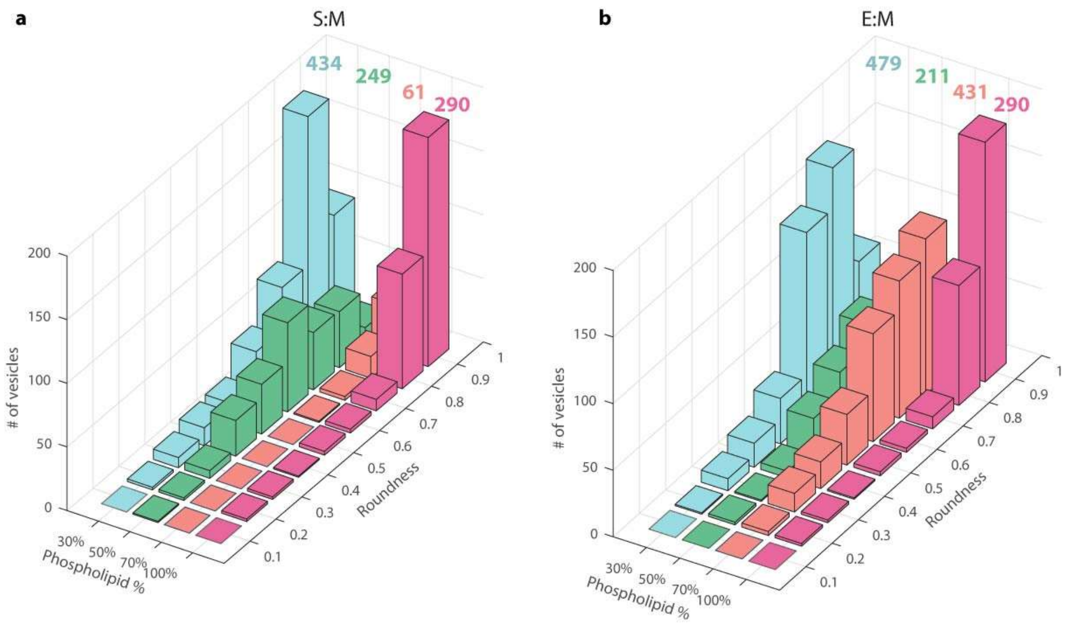


**Fig. S4. Roundness analyses of mixed fatty acid-phospholipid vesicles**. (a) Histograms showing the roundness analyses of mixed Myristoleic acid-Soy bean lipid vesicles. (b) Histograms showing roundness of mixed Myristoleic acid-E-coli lipid vesicles. The percentage of phospholipids are presented in *x* axis and number of vesicles is shown in *y* axis. Each composition is represented with a different color and the number of total analyzed model protocells in each species is stated on top of each set of histograms. *Z* axis shows the roundness values. The analyses of 290 model protocells made from 100% phospholipids containing Soy bean and E-coli lipids are shown both in (a) and (b).

### S5. Free fatty acid quantification

The amount of free fatty acid in lipid solutions was quantified by the Free Fatty Acid Quantification Kit (Sigma-Aldrich) as described by Jin *et al.*[1]. 250 µL of each lipid suspension (with Soy Polar Extract:Myristoleic Acid lipids and Soy Polar Extract:E.Coli Polar Extract 50:49 as a negative control) was prepared as described in the methods. Samples were centrifuged with Accuspin Micro17 (Fischer Scientific, US) for 10 min at 13 500 rpm through AMicon Ultra 3K filters (Merck, Germany). 20 µL of the sample is collected and 5 µL of Fatty Acid Assay Buffer is added. For fluorometric detection, standard suspensions from palmitic acid were prepared with the following concentrations according to the protocol provided by the manufacturer: 0, 0.1, 0.2, 0.3, 0.4, 0.5 nmol/well. Both the standard suspensions and the samples in triplicates were added to a black Cellstar 384-well flat-bottomed plate (Merck/Germany). 2 µL of ACS (Acyl-CoA Synthetase) reagent was then mixed with each sample and the plate was incubated for 30 min at 37 °C. Subsequently, 25 µL of Master mix was prepared with 22 µL Fatty acid Assay Buffer, 1 µL Fatty Acid Probe, 1 µL Enzyme Mix and 1 µL Enhancer Mix, and was added to each sample. The plate was incubated again for 30 min at 37 °C. The fluorescence intensity was measured with a Synergy Neo2 Plate reader (BioTek, US) at an excitation wavelength of 535 nm, and an emission wavelength of 587 nm. An emission curve for the standard suspension of known concentration was obtained and corrected by subtracting the blank value -the reading from the buffer only- from all of the readings. The emission from each measured sample was normalized by subtracting the emission of the negative control which is the sample containing 100% phospholipids and 0% fatty acids (bar in magenta color on the very left). The concentration of the fatty acid in a given sample was calculated as $C = S_a/S_v$ , where $S_a$ was the amount of fatty acid calculated using the standard curve and $S_v$ the sample volume added to the reaction well.

The assay data for two different experiments (two different sets of mixtures) is shown below. The ordinate in **Fig. S5** shows the different percentages of free FA found in each sample.

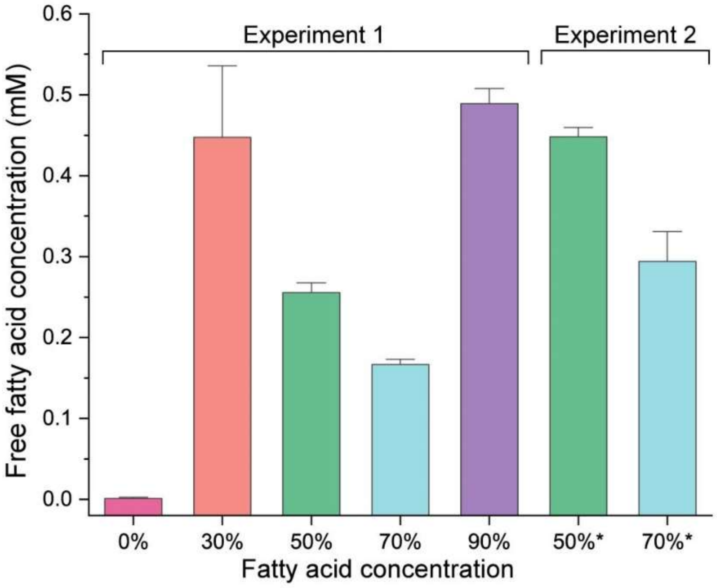


**Figure S5. Free fatty acid quantification**. Histograms show the amount of free fatty acids in solution for various lipid suspensions used in the study. Experiment 2 shows two different suspensions prepared using the same protocol with 50% and 70% FAs. A general trend indicates that the higher the fraction of fatty acid in the vesicle membrane in suspension, the lower the concentration of free fatty acid in solution. 90% FA containing suspension does not follow the trend and has the highest concentration of free FA in solution.

**S6. Encapsulation of fluorescein and RNA inside the protocells (ROIs)**

To expose the surface-adhered protocells to the superfusion media, an open-volume microfluidic pipette (Fluicell AB, Sweden) was used in conjunction with a 3-axis water hydraulic micromanipulator (Narishige, Japan). The superfusion medium consisted of 10 mM HEPES, 100 mM NaCl, 4 mM $CaCl_2$ and 25 µM fluorescein sodium salt (Merck Life Science, Norway), or 25 µM Fl-conjugated 10 base long Poly-Adenine (5'-Fl-AAA AAA AAA A) RNA oligonucleotides (Dharmacon, USA), pH 7.8.

The composition of the lipid mixture and superfusion medium is noted on the upper left corner of the micrograph (*flu* for fluorescein, and *RNA*). Selected regions of interest (ROIs) are shown as circles in dashed lines. From each experiment, 7 lipid compartments were chosen. One ROI was placed inside each compartment, and one besides it in order to capture the fluorescence intensity of the background. Compartments marked with '*x*' in the micrographs either relocate during the recording, or collapse during the experiment hence were not chosen as ROIs. The fluorescence intensities in the selected ROIs were used to obtain the graphs in **Fig. 2h** and **i** of the main manuscript. Scale bar: 10 µm.

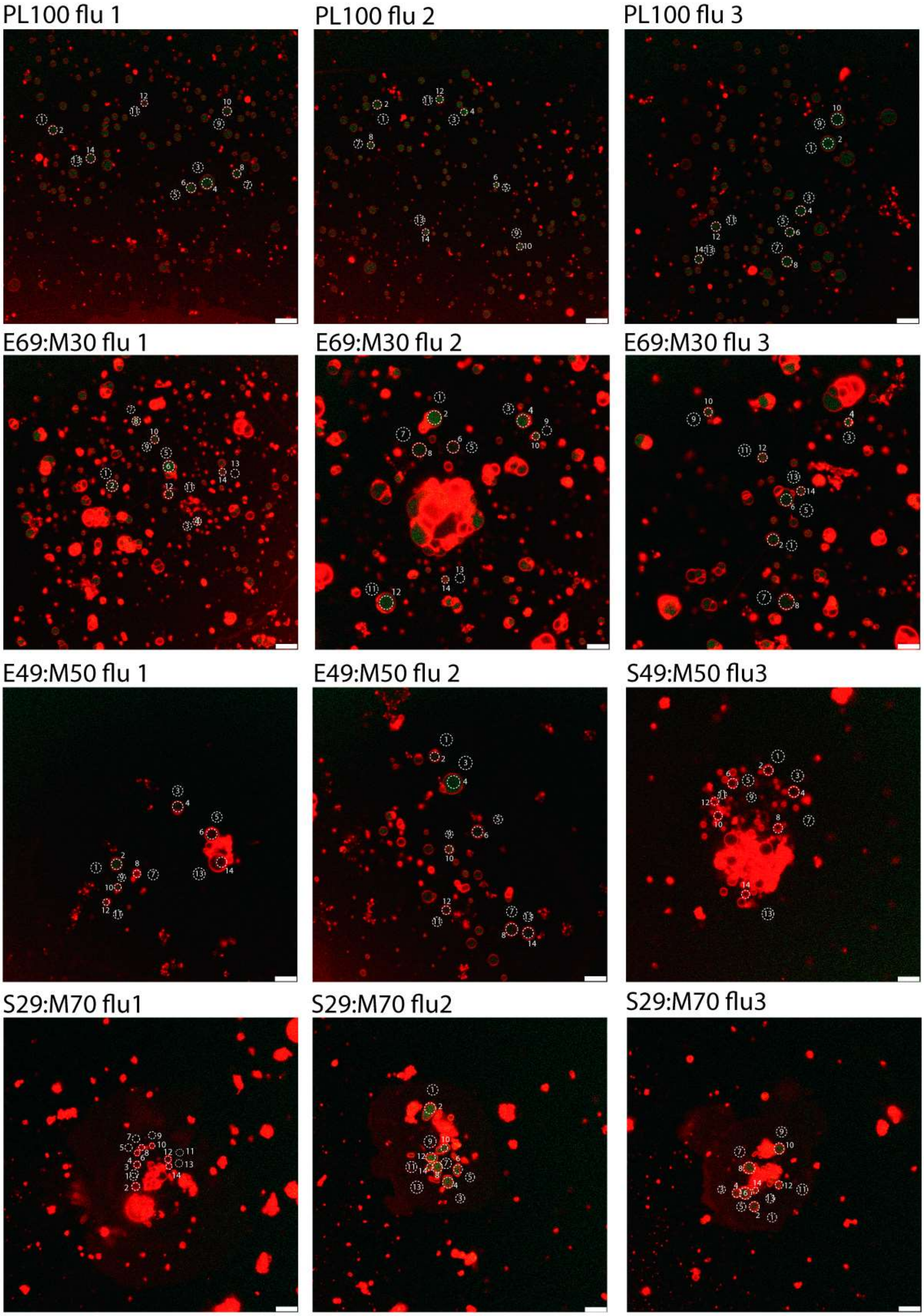
PL100 flu 1
PL100 flu 2
PL100 flu 3
E69:M30 flu 1
E69:M30 flu 2
E69:M30 flu 3
E49:M50 flu 1
E49:M50 flu 2
S49:M50 flu3
S29:M70 flu1
S29:M70 flu2
S29:M70 flu3

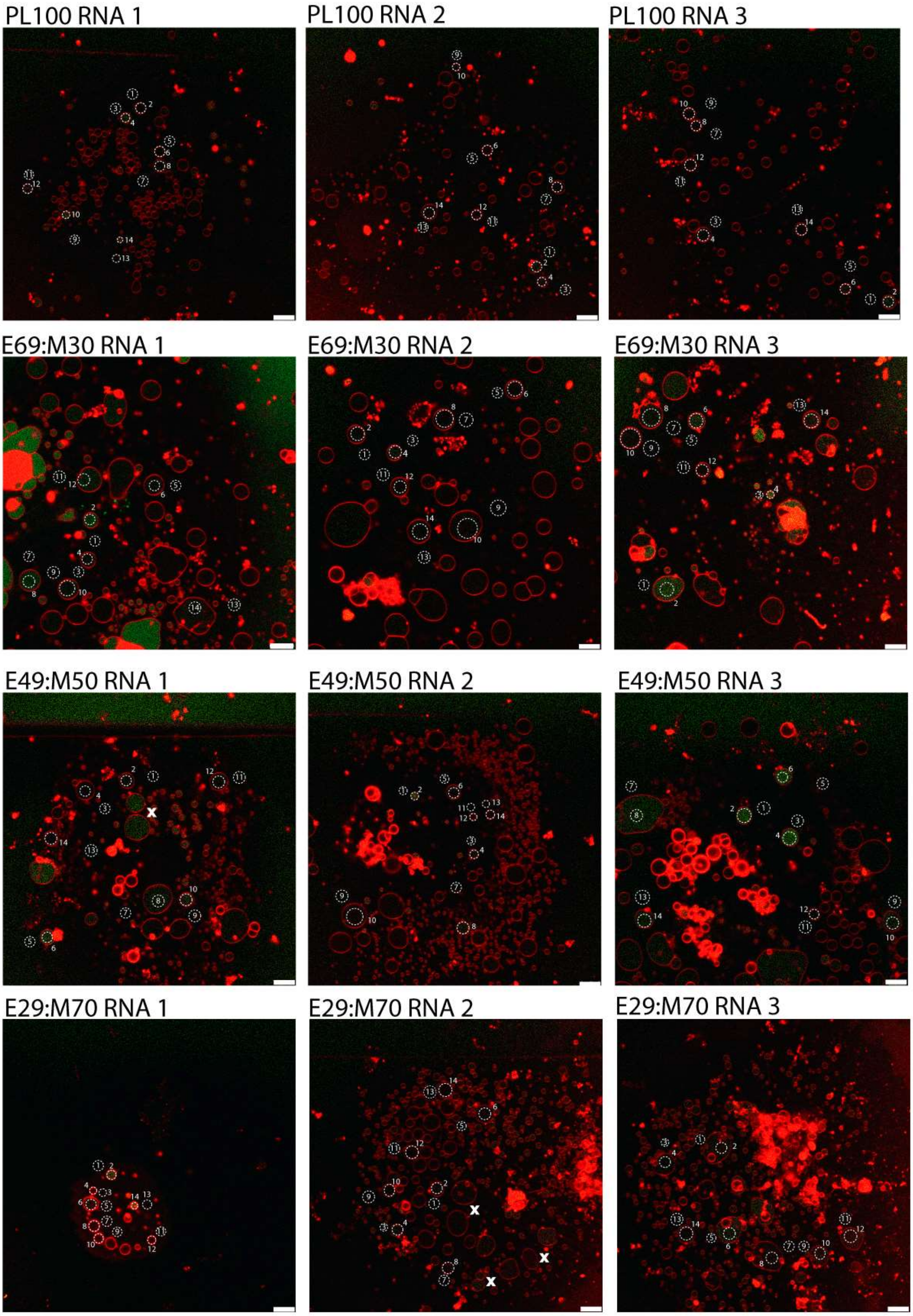
PL100 RNA 1
PL100 RNA 2
PL100 RNA 3
E69:M30 RNA 1
E69:M30 RNA 2
E69:M30 RNA 3
E49:M50 RNA 1
E49:M50 RNA 2
E49:M50 RNA 3
E29:M70 RNA 1
E29:M70 RNA 2
E29:M70 RNA 3

### S7. Movie captions

**Movie S1. Uptake of RNA by model protocells from 70PL:30FA.** Movie S1 shows an example of RNA uptake in a region shown in 'E69:M30 RNA 3' in **SI 6**. At the start of the movie, the superimposed transmission channel shows the position of the pipette tip. Later it is removed to show the encapsulation of media by the compartments only in the fluorescence emission channels. Red and green (false) colors represent rhodamine-fluorophore conjugated membrane and fluorescein-conjugated RNA, respectively. Sample ROIs are represented as dashed lines.
